## Supplementary Text and Figures for "Developmental hematopoietic stem cell variation explains clonal hematopoiesis later in life"

### Supplementary Information for
#### Developmental hematopoietic stem cell variation explains clonal hematopoiesis later in life

Jesse Kreger[a], Jazlyn A. Mooney[a], Darryl Shibata[b], Adam L. MacLean[a,*]

[a]Department of Quantitative and Computational Biology, University of Southern California, Los Angeles, CA, USA

[b]Department of Pathology, Keck School of Medicine, University of Southern California, Los Angeles, CA, USA

---

#### Supplementary Text

**S1 Selection criteria and full list of "neutral" fCpG sites** **2**

**S2 Comparison to population level fluctuating methylation clock models** **2**

**S3 Cell replacement and methylation rates define range of FMC model behavior** **3**

**S4 Cell threshold at which the embryos split and number of cell clones effect clonal variation between twins** **3**

**S5 Further details on selecting $N_{\text{split}}$ for simulations of twins** **4**

**S6 Variants with weak selection explain FMC dynamics for dizygotic twins and unrelated individuals** **4**

#### Supplementary Figures

S1 **Effect of varying cell replacement rate $\alpha$** . . . . . . . . . . . . . . . . . . . . 6
S2 **Effect of varying (de)methylation rate $\gamma$** . . . . . . . . . . . . . . . . . . . 7
S3 **Effect of varying number of cell clones $N_{\text{clones}}$.** . . . . . . . . . . . . . . . . . 8
S4 **Determining $N_{\text{split}}$ for monozygotic and dizygotic twins** . . . . . . . . . . 9
S5 **Distribution of initial Pearson coefficient** . . . . . . . . . . . . . . . . . . . 10
S6 **Variants with weak selection arise during development and explain FMC dynamics for dizygotic twins** . . . . . . . . . . . . . . . . . . . . . . . . . 11
S7 **Variants with weak selection arise during development and explain FMC dynamics for unrelated individuals** . . . . . . . . . . . . . . . . . . . . . 12
S8 **Model vs data comparison favors no or weak selection** . . . . . . . . . . . . 13

#### Supplementary Tables

S1 **Example of percent fitness advantage for each selection regime** . . . . . . . 14
S2 **Model vs data comparison metric** . . . . . . . . . . . . . . . . . . . . . . . 14

---

### Supplementary Text

#### S1 Selection criteria and full list of "neutral" fCpG sites

Similar to[1], we study the dynamics of fCpG sites that are "neutral", i.e. CpG loci that are not actively regulated and that show a high degree of intraindividaul heterogeneity. To filter out "neutral" fCpG sites, we implement the following filtering pipeline using dataset GSE40279[2]:

1. average methylation for each probe from provided GSE40279[2] beta file with 656 normal blood samples
2. select CpG loci with mean values between 0.4 and 0.6
3. select CpG loci with higher variance between individuals
4. remove likely single nucleotide polymorphism based on consistent outlier values ($> 0.85$, $< 0.15$) in individual blood samples
5. remove more cell type specific methylation based on purified blood cell types (granulocytes, CD4+, CD8+, CD56+, monocytes, B cells, PBMCs, eosinophils, data from GSE35069[3]) with skewed average methylation ($> 0.6$, $< 0.4$)

A full list of the 3,918 "neutral" fCpG sites used in this study can be found here: github.com/maclean-lab/scFMC-model.

#### S2 Comparison to population level fluctuating methylation clock models

Our model is built to allow for single-site single-cell resolution of FMCs. At the population level, it is similar to previous FMCs models that allow for analysis of average methylation profiles in a population of cells, such as in[1,4,5].

As noted in the main text, in[1] the authors develop a stochastic differential equation mathematical model to measure human adult stem cell dynamics from methylation arrays. Their study is the basis for our model. Some key differences are that the model developed in[1] assumes:

- a population model, where the state of their system of $N_{\text{cells}}$ cells can be fully characterized using just two state variables (the number of stem cells containing a single methylated allele and the number of stem cells containing two methylated alleles). In contrast, we develop a model allowing for the analysis of single-site single-cell resolution of FMCs (which comes at a computational cost).
- that cell replacement and (de)methylation are independent, i.e. that a fCpG site can flip-flop in the absence of a cell replacement event. We assume that (de)methylation events occur during cell replacement.
- analysis of human adult stem cells. Our model importantly includes embryo development, and how variation in development can define HSC dynamics later in life.

At the population level, the models are similar, and we also show that for healthy individuals fluctuating methylation revealed unimodal distributions centered around 50% methylation.

In[4] phylogenetic trees and sequencing of genomes from single cell-derived colonies of haematopoietic cells are used to show that haematopoiesis in elderly humans shows significantly decreased clonal diversity. They simulate a phylogenetic model and show that constant acquisition of driver mutations with moderate fitness benefits entirely explained the abrupt change in clonal diversity.

Our methylation data (Fig. 1C-F in the main text) and simulations of our model (Fig. 4-5 in the main text) show similar loss of clonal diversity after 60 years of age. However, we suggest that moderate fitness effects are not needed to explain the data, instead weak selection with variants in development result in the similar dynamics.

In[5] coalescent theory is used to estimate the net growth rate of clones from either reconstructed phylogenies or the number of shared mutations. The authors show that cell clones with multiple driver mutations have significantly increased growth rates, which are associated with shorter time to disease diagnosis. In contrast, our model represents an evolutionary approach using FMCs to measure clonal diversity and methylation profiles. While experimentally continuous observation of clonal architecture in cancer evolution is impossible, simulations of our model provide single-site single-cell resolution that are not possible in the clinic.

### S3 Cell replacement and methylation rates define range of FMC model behavior

The two main events in the model are cell replacement (with rate $\alpha$) and (de)methylation (with rate $\gamma$), full details are described in main text Methods. As described in main text Table 2, for simulation of the model we generally use $\alpha \approx \frac{1}{365}$ based on[4,6] and $\gamma \approx 10^{-3}$ based on[1,7]. These two parameters define model dynamics, in particular,

- If $\alpha$ is too low, then nothing happens as the population of cells will remain relatively unchanged over time. If $\alpha$ is too high, then cell turnover happens too frequently and stochastic fixation will happen even in the case of little or no fitness differences ($s_i = 0$ for all $i = 1, 2, \ldots, N_{\text{clones}}$) which does not match what we see in the methylation data (see Methods in the main text). These trends can be seen in Fig. S1. Fig. S1A shows how a large rate of cell replacement can create a rapid change in clonal dynamics and cause domination of a few cell clones even in the case of small or no fitness differences. Fig. S1A (middle panel) shows how the Pearson coefficient can rapidly decrease in the case that different cell clones rise to dominance in the two individuals. Fig. S1B shows model dynamics with a reasonable cell replacement rate[4,6]. Fig. S1C shows how a small rate of cell replacement results in stagnant dynamics, as the population of cells does not change much over time.
- If $\gamma$ is too low, then the dynamics are completely determined by cell replacement (which does not match fCpG data/site selection procedure, see Section S1). If $\gamma$ is too high, then the methylation distribution will be unimodal (and centered at $\frac{1}{2}$) but will be too tightly distributed (with too small variance) which does not match the methylation data (see Methods in the main text). In particular, as $\gamma$ increases the average methylation distribution goes from bimodal peaks at 0 and 1, to a trimodal W distribution, to a unimodal distribution with a peak at $\frac{1}{2}$ (see also Fig. 3A in[1]). These trends can be seen in Fig. S2. Fig. S2A shows how a large rate of (de)methylation results in very narrow methylation distributions (Fig. S2A right panel, 100 years) which is not seen in the methylation data. Fig. S1B shows model dynamics with a reasonable (de)methylation rate[1,7]. Fig. S2C shows that if $\gamma$ is too low, then the dynamics are completely determined by cell replacement (which also does not match the data) and results in very wide unimodal methylation distributions (Fig. S2C right panel, 100 years).

### S4 Cell threshold at which the embryos split and number of cell clones effect clonal variation between twins

Here we analyze the effect of the cell threshold at which the embryos split, $N_{\text{split}}$ (Fig. 2C in the main text) and number of cell clones, $N_{\text{clones}}$ (Fig. S3) when there is variation during development (frequency-dependent growth, see main text Methods for model description). For both figures the white striped bars represent the clonal distribution in the shared embryo when it reaches $N_{\text{split}}$ cells and splits into twins. Yellow bars represent individual 1 and red bars represent individual 2 when the embryos have reached $N_{\text{cells}}$ cells and are finished growing.

Fig. 2D-E in the main text shows the effect of increasing the cell threshold at which the embryos split $N_{\text{split}}$. Here, we see that for a lower splitting threshold $N_{\text{split}} = 10$, the clonal distributions of the twins (yellow vs red bars) are extremely different, and are also very different from the clonal distribution of the shared embryo (white bars bars). This is because the earlier the embryos splits, the higher likelihood there is for variation in the twins and for different clones to rise to prominence. In the case of a higher splitting threshold $N_{\text{split}} = 500$, the two twins clonal distributions at birth (when the embryos have reached $N_{\text{cells}}$ cells) are much more similar. This will also result in a higher initial Pearson coefficient (see Section S5 and Fig. S4).

Fig. S3 shows the effect of increasing the number of cell clones $N_{\text{clones}}$. Here, we see that for a lower number of cell clones $N_{\text{clones}} = 3$ (Fig. S3A), the clonal distributions of the twins (yellow vs red bars) are more similar than for a higher number of cell clones ($N_{\text{clones}} = 5$, Fig. S3B). This is because fewer cell clones results in less variability in the clonal distributions, as well as a higher initial Pearson coefficient.

#### S5 Further details on selecting $N_{\text{split}}$ for simulations of twins

As described in Methods in the main text (see also Fig. 2C), the value of $N_{\text{split}}$ is determined based on simulating the model and finding the best fit to Pearson coefficients at birth from methylation data from twins. Fig. S4 shows the data from Fig. 2C in the main text with 10 cell clones (Fig. 2C green data points). Here, we use the LsqFit.jl Julia curve fitting package[8] to fit this data to a Hill-type function (black line, Fig. S4) of the form

$$f(x) = \frac{p_1 x}{x + p_2} \tag{S1}$$

where $p_1$, $p_2$ are parameters, $x$ is the variable representing $N_{\text{split}}$, and $f(x)$ represents the Pearson coefficient between twins at birth. For monozygotic twins (initial Pearson coefficient 0.87) we find that $N_{\text{split}} \approx 36$, for dizygotic twins (initial Pearson coefficient 0.72) $N_{\text{split}} \approx 15$, and for unrelated individuals (initial Pearson coefficient 0.61) $N_{\text{split}} \approx 10$. Fig. S5 shows an example distribution of the initial Pearson coefficient for $2 \times 10^2$ simulations with $N_{\text{clones}} = 10$ and $N_{\text{split}} = 36$.

#### S6 Variants with weak selection explain FMC dynamics for dizygotic twins and unrelated individuals

Fig. 4 in the main text shows variants with weak selection (see Table S1) that arise during development and explain FMC dynamics for monozygotic twins. Here we show similar results for dizyogitc twins and unrelated indiviuals. In particular, Fig. S6 shows simulations for dizygotic twins and Fig. S7 shows simulations for unrelated individuals as in Fig. 4 in the main text.

Fig. S8 shows comparisons of simulated model trajectories (gray lines) with simulated data trajectories (colored lines). Table S2 shows the the mean distance from each model simulation to each data trajectory, see main text Methods for data trajectory details.

### Supplementary Figures

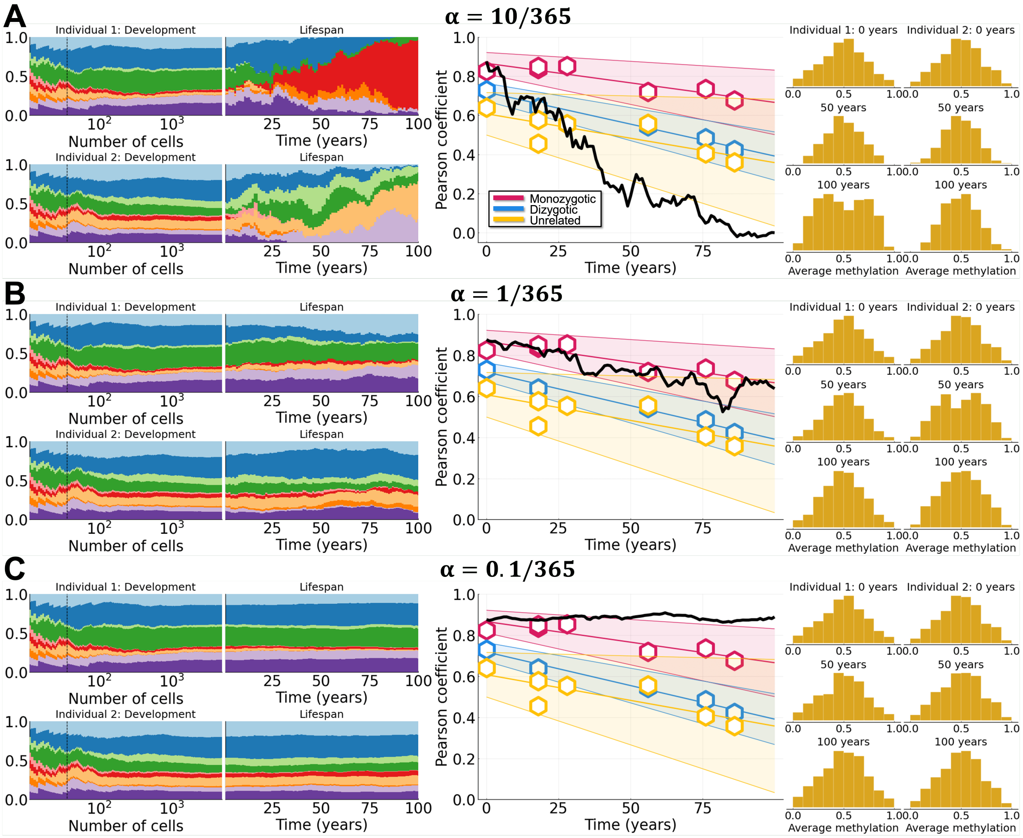


Figure S1: **Effect of varying cell replacement rate** $\alpha$**.** Effect of varying the cell replacement rate with weak selection ($a = 0.05,\ \theta = 0.01$). Simulations shown for monozygotic twins ($N_{\text{split}} = 36$) with variation (frequency-dependent growth) during development. For Pearson correlation plots hexagons represent means of datasets and 90% confidence intervals are shown. **A-C**: Clone growth frequency plots for both individuals during development and life (dashed vertical line represents $N_{\text{split}}$), Pearson correlation coefficient, and $\beta$ distributions at 0, 50, and 100 years of life. A: Large cell replacement rate ($\alpha = \frac{10}{365}$). B: Medium cell replacement rate ($\alpha = \frac{1}{365}$). C: Small cell replacement rate ($\alpha = \frac{0.1}{365}$). All other parameter values can be found in Table 2 in the main text.

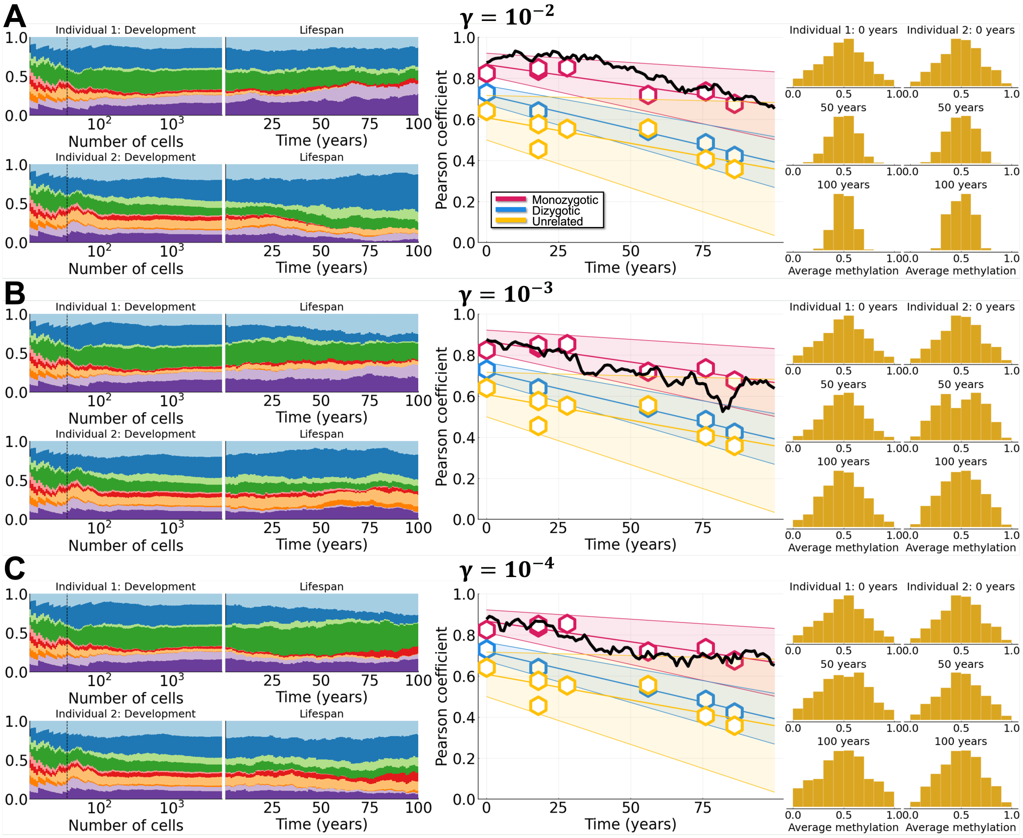


Figure S2: **Effect of varying (de)methylation rate** $\gamma$**.** Effect of varying the (de)methylation rate with weak selection ($a = 0.05,\ \theta = 0.01$). Simulations shown for monozygotic twins ($N_{\text{split}} = 36$) with variation (frequency-dependent growth) during development. For Pearson correlation plots hexagons represent means of datasets and 90% confidence intervals are shown. **A-C**: Clone growth frequency plots for both individuals during development and life (dashed vertical line represents $N_{\text{split}}$), Pearson correlation coefficient, and $\beta$ distributions at 0, 50, and 100 years of life. A: Large (de)methylation rate ($\gamma = 10^{-2}$). B: Medium (de)methylation rate ($\gamma = 10^{-3}$). C: Small (de)methylation rate ($\gamma = 10^{-4}$). All other parameter values can be found in Table 2 in the main text.

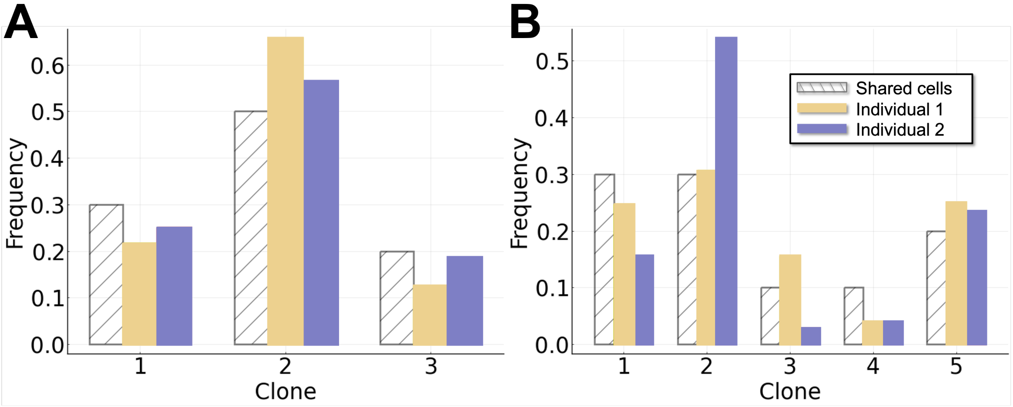


Figure S3: **Effect of varying number of cell clones $N_{\mathbf{clones}}$.** White striped bars represent the clonal distribution in the shared embryo when it reaches $N_{\text{split}}$ cells and splits into two embryos. Yellow/blue bars represent individuals 1 or 2 at the point at which the embryo has reached $N_{\text{cells}}$ cells and finished growing. Here we use $N_{\text{split}} = 10$, weak selection, and assume variation during development. A: $N_{\text{clones}} = 3$. B: $N_{\text{clones}} = 5$.

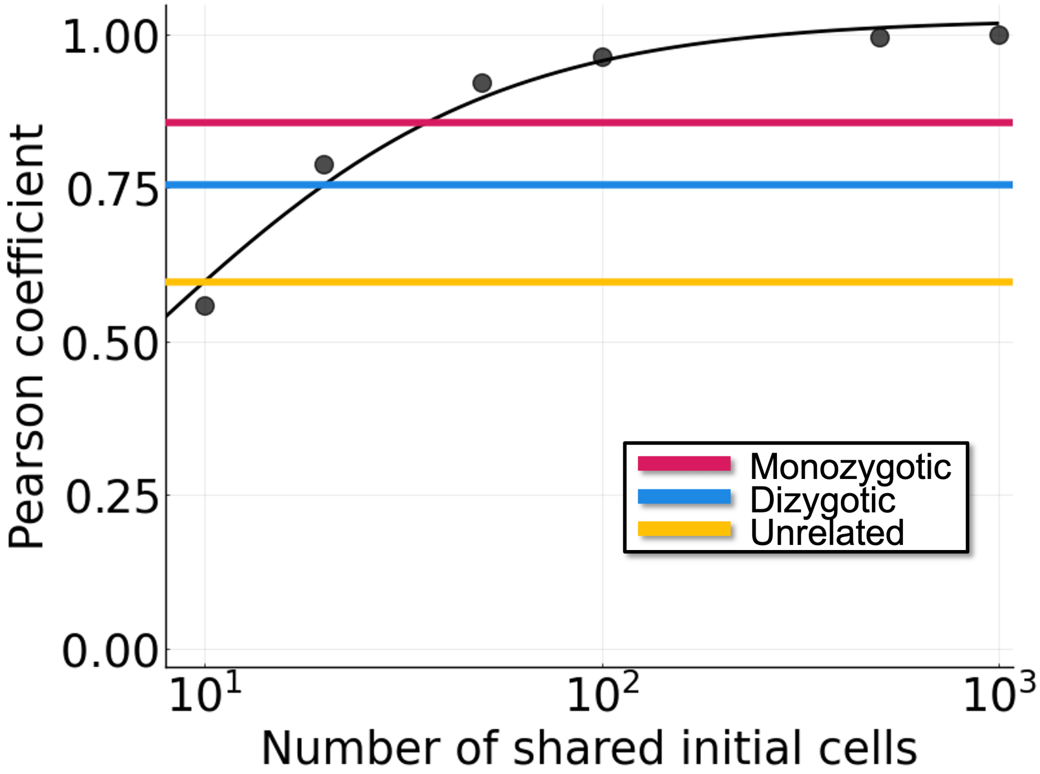


Figure S4: **Determining $N_{\mathbf{split}}$ for monozygotic and dizygotic twins.** The value of $N_{\mathrm{split}}$ is determined based on simulating the model and finding the best fit to Pearson coefficients at birth from methylation data from twins. The black data points represent the data from Fig. 2C in the main text with 10 cell clones (Fig. 2C green data points). We use a Hill-type function (black line, Equation S1) to approximate $N_{\mathrm{split}}$. The horizontal lines represent the initial Pearson coefficient at birth for monozygotic twins (red), dizygotic twins (blue) and unrelated individuals (yellow). For monozygotic twins (initial Pearson coefficient 0.85) we find that $N_{\mathrm{split}} = 36.11$, for dizygotic twins (initial Pearson coefficient 0.75) $N_{\mathrm{split}} = 19.98$, and for unrelated individuals (initial Pearson coefficient 0.60) $N_{\mathrm{split}} = 10.00$.

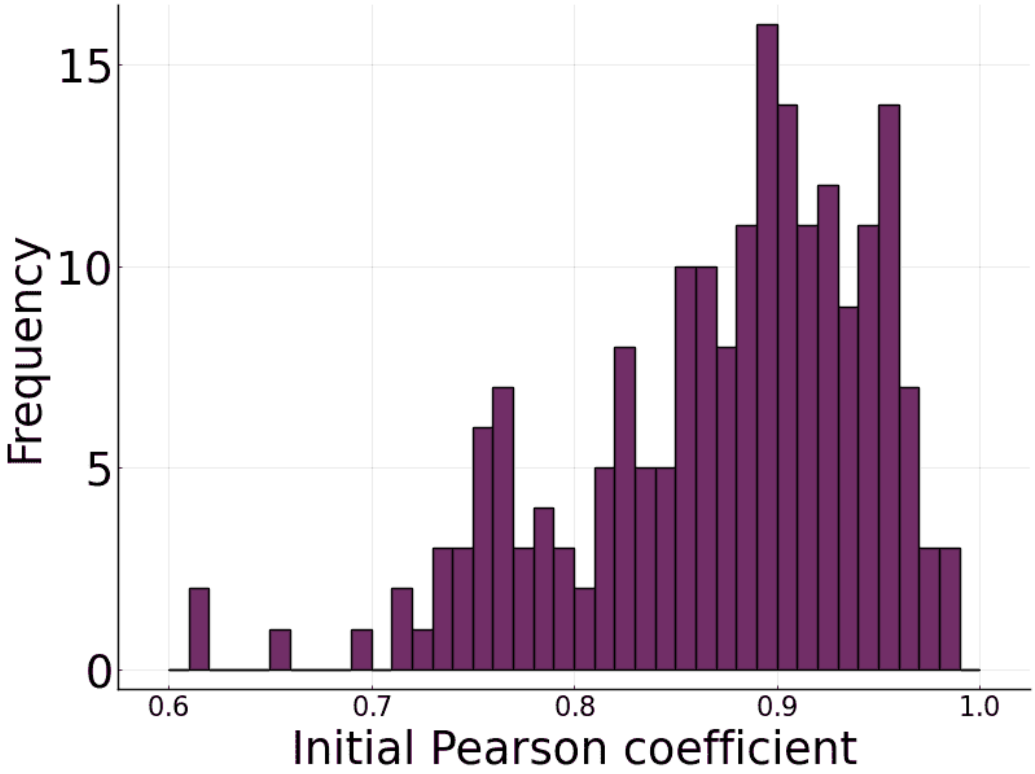


Figure S5: **Distribution of initial Pearson coefficient.** Histogram representing the distribution of the initial Pearson coefficient for $2 \times 10^2$ simulations. Here we assume $N_{\mathrm{clones}} = 10$, $N_{\mathrm{split}} = 36$, and variation during development with weak selection. The mean initial Pearson coefficient is 0.87 and the standard deviation is 0.074.

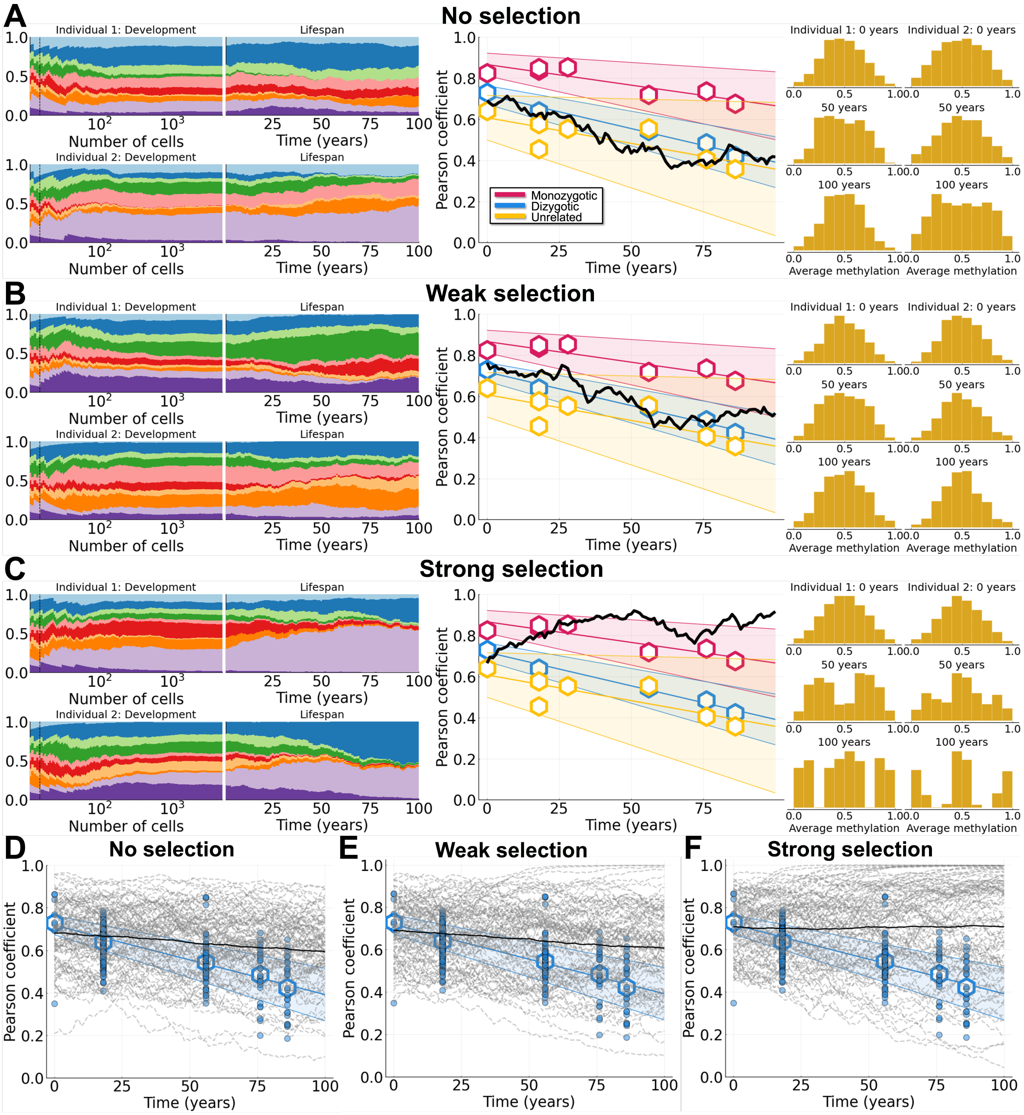


Figure S6: **Variants with weak selection arise during development and explain FMC dynamics for dizygotic twins.** Simulations shown for dizygotic twins ($N_{\text{split}} = 15$) with variation (frequency-dependent growth) during development. For plots showing Pearson correlation data (middle column and bottom row), dots represent individual comparisons, hexagons represent means of datasets, and shaded bands represent 90% confidence intervals. **A-C**: Clone growth frequency plots for both individuals during development and life (dashed vertical line represents $N_{\text{split}}$), Pearson correlation coefficient, and $\beta$ distributions at 0, 50, and 100 years of life. A: No selection. B: Weak selection ($a = 0.05$ and $\theta = 0.01$). C: Strong selection ($a = 0.05$ and $\theta = 0.05$). **D-F**: Results from $10^2$ simulations are shown (dashed lines are individual simulations and solid lines are mean trajectories). D: No selection. E: Weak selection ($a = 0.05$ and $\theta = 0.01$). F: Strong selection ($a = 0.05$ and $\theta = 0.05$). All other parameter values can be found in Table 2 in the main text.

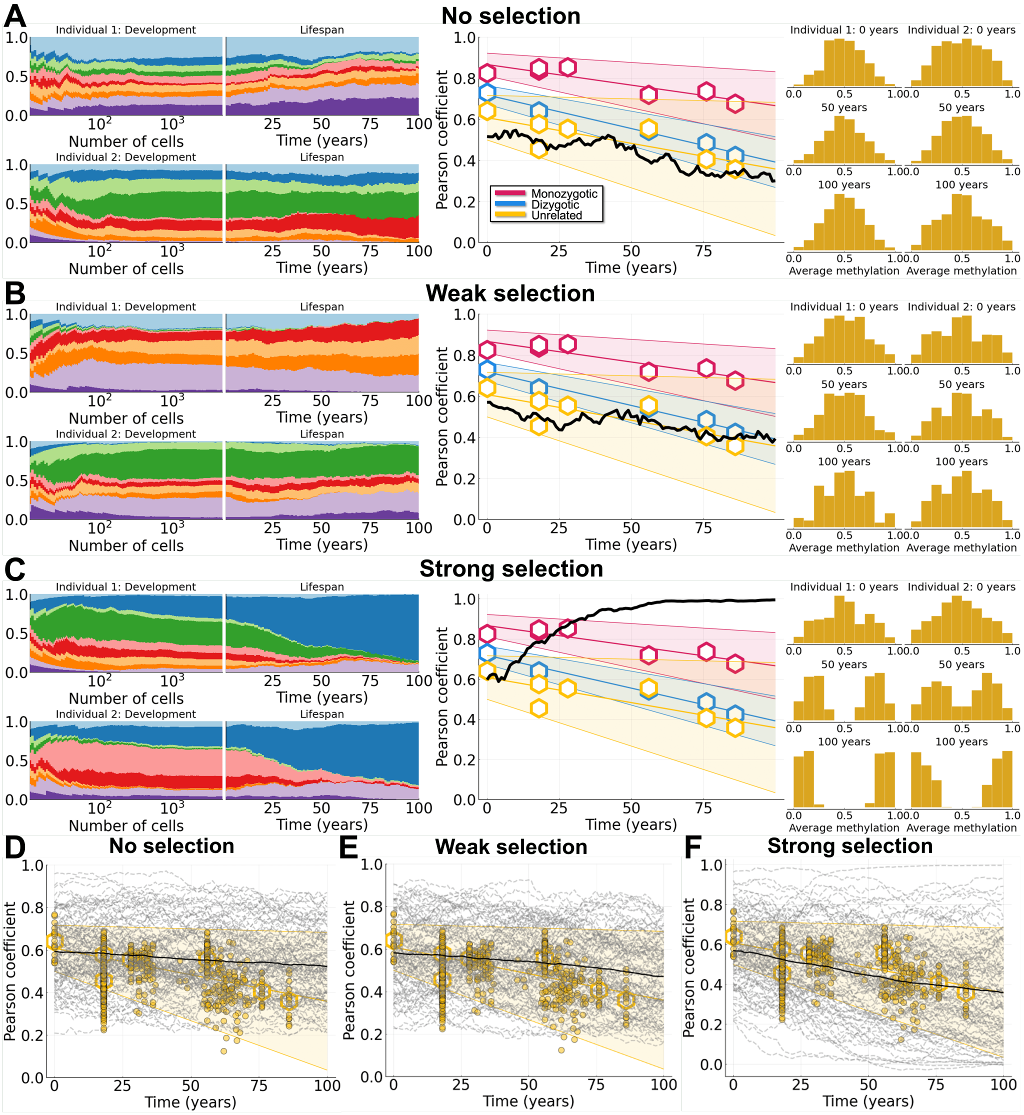


Figure S7: **Variants with weak selection arise during development and explain FMC dynamics for unrelated individuals.** Simulations shown for unrelated individuals ($N_{\text{split}} = 10$) with variation (frequency-dependent growth) during development. For plots showing Pearson correlation data (middle column and bottom row), dots represent individual comparisons, hexagons represent means of datasets, and shaded bands represent 90% confidence intervals. **A-C**: Clone growth frequency plots for both individuals during development and life (dashed vertical line represents $N_{\text{split}}$), Pearson correlation coefficient, and $\beta$ distributions at 0, 50, and 100 years of life. A: No selection. B: Weak selection ($a = 0.05$ and $\theta = 0.01$). C: Strong selection ($a = 0.05$ and $\theta = 0.05$). **D-F**: Results from $10^2$ simulations are shown (dashed lines are individual simulations and solid lines are mean trajectories). D: No selection. E: Weak selection ($a = 0.05$ and $\theta = 0.01$). F: Strong selection ($a = 0.05$ and $\theta = 0.05$). All other parameter values can be found in Table 2 in the main text.

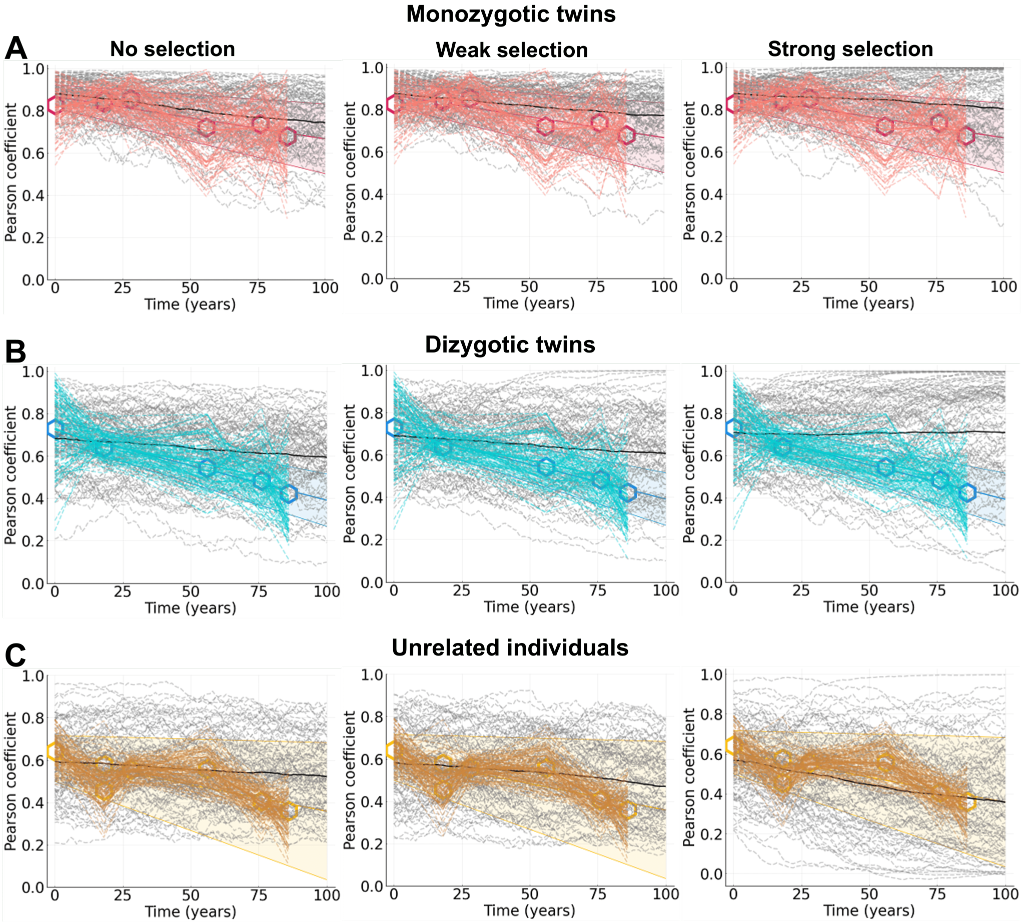


Figure S8: **Model vs data comparison favors no or weak selection.** $10^2$ simulations shown for monozygotic twins ($N_{\text{split}} = 36$), dizygotic twins ($N_{\text{split}} = 15$), and unrelated individuals ($N_{\text{split}} = 10$) with variation (frequency-dependent growth) during development. Left panels shown no selection, middle panels show weak selection, and right panels show strong selection. Hexagons represent means of datasets and 90% confidence intervals are shown. Model simulations are in gray and data trajectories (see Methods for details) are in red (monozygotic twins), blue (dizygotic twins), and yellow (unrelated individuals). A: Monozygotic twins. B: Dizygotic twins. C: Unrelated individuals. All other parameter values can be found in Table 2 in the main text.

### Supplementary Tables

| Clone | No selection | Weak selection ($a = 0.05,\ \theta = 0.01$) | Strong selection ($a = 0.05,\ \theta = 0.05$) |
|---|---|---|---|
| 1 | 0.0 | 0.032 | 0.083 |
| 2 | 0.0 | $7.2 \times 10^{-8}$ | $2.0 \times 10^{-7}$ |
| 3 | 0.0 | $1.5 \times 10^{-9}$ | 0.0 |
| 4 | 0.0 | $2.6 \times 10^{-3}$ | 0.0 |
| 5 | 0.0 | 0.0 | 3.3 |
| 6 | 0.0 | $9.7 \times 10^{-5}$ | 0.0 |
| 7 | 0.0 | 0.0 | 0.036 |
| 8 | 0.0 | **0.25** | 0.0 |
| 9 | 0.0 | $3.4 \times 10^{-9}$ | **6.2** |
| 10 | 0.0 | $5.7 \times 10^{-7}$ | 0.0 |

Table S1: **Example of percent fitness advantage for each selection regime.** Example of coefficients ($s_i$), chosen from a Gamma distribution with $a = 0.05,\ \theta = 0.01$ (weak selection) or $a = 0.05,\ \theta = 0.05$ (strong selection). The most fit clone (largest fitness coefficient) is in bold for each case. Selection coefficients $s_i < 10^{-10}$ are rounded to 0.

| | No selection | Weak selection ($a = 0.05,\ \theta = 0.01$) | Strong selection ($a = 0.05,\ \theta = 0.05$) |
|---|---|---|---|
| **Monozygotic** | 0.134 | 0.132 | 0.143 |
| **Dizygotic** | 0.182 | 0.191 | 0.231 |
| **Unrelated individuals** | 0.163 | 0.160 | 0.186 |

Table S2: **Model vs data comparison metric.** To directly compare model simulations to the methylation data we compute the function $d$, which measures the mean distance from each model simulation to each data trajectory at the given time points. See main text Methods for data trajectory details.
